# The integrative ecology of the saddleback anemonefish, *Amphiprion polymnus*, reveals it is a host generalist across multiple ecological axes

**DOI:** 10.64898/2026.06.02.729679

**Authors:** Catheline Y.M. Froehlich, Benjamin M. Titus

## Abstract

Understanding how symbiosis generates biodiversity requires comprehensive insight into the underlying ecology to reveal broad evolutionary patterns, and the clownfish-anemone symbiosis serves as an excellent model. Recently, a new paradigm revealed sea anemone host associations of adult clownfish drove convergent evolution of host specialists and generalists. An alternative study downplayed host specificity in explaining clownfish evolution due partly to an incongruence between the behavior, biology, and host specificity of a single species—*Amphiprion polymnus*. Previously classified as a host specialist, we revisit the host specificity and ecology of *A. polymnus* by studying its host associations, behavioral, and dietary ecology compared to a co-occurring host generalist and specialist. By incorporating n = 358 observations, we find *A. polymnus* associates with five sea anemone species, including a newly documented host, and four at the adult life stage. We show that *A. polymnus* swim consistently far from their hosts and have broad dietary niches, an ecology that aligns with generalists. Cumulatively, our data indicate *A. polymnus* is a host generalist, and not a specialist as previously classified. Our findings serve to reconcile previously conflicting studies about *A. polymnus*, and reinforce the hypothesis that host is the key ecological variable for understanding clownfish evolution.

**Suitable subject areas:** behaviour, ecology, evolution

## 1. Introduction

Symbioses are tightly linked ecological interactions that persist over evolutionary timescales (e.g. [1–3]). An evolutionary understanding of how symbiosis generates and maintains biodiversity thus requires a synthetic understanding of the underlying ecology of the interaction. Commonly, ecological information within symbioses is synthesized by quantifying the number of ecological associations between symbiotic organisms, thereby creating association matrices and classifying species as specialists or generalists based on raw association counts (e.g. [4–7]). Yet this measure alone may not fully capture the most ecologically and biologically relevant aspects of the interaction. Of critical importance is to properly characterize the strength and frequencies of the ecological interaction (a.k.a. specificity) alongside other ecological variables linked to the interaction, e.g. behaviors, trophic positions, and phenotypic evolution, e.g. life form, body size, color patterns, defenses [8–12]. Synthesizing all ecological aspects of a symbiosis and integrating them into a well-resolved evolutionary framework is a more powerful approach for understanding symbiosis as an important driver of global biodiversity.

The adaptive radiation of clownfishes is a relevant case that demonstrates both the futility of using inadequate association counts, and utility of more ecologically meaningful metrics, to understand the role of symbiosis in generating biodiversity [5,8,13,14]. Clownfishes, or anemonefishes, are a clade of 29 species in the genus *Amphiprion* that form obligate mutualisms with host sea anemones (Cnidaria: Hexacorallia: Actiniaria)[15]. Their adaptation to live with venomous sea anemones is held as the key ecological innovation and driver of the adaptive radiation of the genus [5,8,13,14,16,17]. However, early phylogenetics work using raw association counts failed to recover an ecological and phylogenetic signal that could broadly explain diversification in the genus beyond geography [5,13,14,16]. Recently, Gaboriau et al. [8] revised the clownfish host association matrix, discovering 29 new host associations across 10 clownfish species. Gaboriau et al. [8] quantified host association frequencies (i.e. number of observations that a fish was found in a host species). Importantly, Gaboriau et al. [8] hypothesized that the species and frequency with which clownfishes use different sea anemones in their adult life stage (i.e. reproductive stage) captures the most ecologically and biologically relevant aspects of the symbiosis. Gaboriau et al. [8] called these “reproductive host associations” and found that host use has driven the evolution of convergent clownfish color patterns, morphology, and genomes. Three distinct color patterns evolved based on reproductive host associations[8]: orange fish living on *Radianthus magnifica,* red fish living on *Entacmaea quadricolor*, and black and white fish living on many hosts (i.e. host generalist). Ultimately, reproductive host association is the first ecological variable with a significant signal able to broadly explain clownfish diversification and color pattern evolution [8].

Prior to Gaboriau et al. [8] there was a rich body of literature exploring the evolution and function of clownfish color patterns (e.g. [18–25]). These demonstrated secondary color-pattern functions related to intra- and inter-specific social signaling yet could not comprehensively explain the evolution of color pattern across the entire clade. With host specificity now identified as the ultimate evolutionary driver of convergent clownfish color patterns and morphology, there has been increased attention into the underlying ecology, function, and genomics of their iconic color patterns because their convergence implies shared selective pressures [8,26,27]. Building on Gaboriau et al. [8], Froehlich et al. [26] used an integrative approach to show that host specialists have different underlying ecologies than generalists. Key findings [26] indicate that both categories of host specialists swim close to their host, background match to their anemones, and have narrow dietary niches and microbiomes. In contrast, host generalists swim far from their host, are highly visible against their anemones, and have broader dietary niches and microbiomes [26]. While not all 29 clownfish species fit neatly into these three reproductive host use and color pattern categories, a cohesive narrative is emerging that sea anemone use has broadly driven the biodiversity, ecology, and phenotypic evolution of clownfishes.

A recent alternative hypothesis by Mercader et al. [28] however argues that clownfish eco-morphotypes evolved independently of host specificity. Mercader et al. [28] studied the behavior, swimming ability, physiology, and musculature of six clownfish species: *A. perideraion, A. ocellaris*, *A. frenatus*, *A. clarkii*, *A. polymnus,* and *A. sandaracinos*). Mercader et al. [28] conclude that clownfish phenotypic evolution is decoupled from host specificity rest in part on the similarities recovered between *A. clarkii* and the saddleback anemonefish, *A. polymnus* [28]. Although distantly related, both fish have black body colors with multiple thick white bars, similar behaviors, swimming abilities, physiology, and musculature, yet differ in their host specificities: *A. clarkii* is a host generalist while *A. polymnus* is a *Stichodactyla* specialist [8,28]. *Amphiprion polymnus* is thus an outlier to the clownfish color-pattern and host use classification scheme recovered by Gaboriau et al. [8] and important to the alternative hypothesis proposed by Mercader et al [28], downplaying the role of the host anemones as the primary driver of the diversification of clownfishes.

A synthetic and comprehensive understanding of the ecology of the symbiosis between *A. polymnus* and their host anemones is thus important for continuing to evaluate alternative hypotheses proposed by Gaboriau et al. [8] and Mercader et al. [28] on the diversification of clownfishes. In their previous classification scheme, Gaboriau et al. [8] documented and verified *A. polymnus* in association with three species of host sea anemones (*S. haddoni*, *Radianthus doreensis,* and *Heteractis aurora*), with only two sea anemones serving as adult reproductive hosts (*S. haddoni*, *R. doreensis*). As 88% of all reproductive adult associations for *A. polymnus* occurred with Haddon’s carpet anemone *S. haddoni*, Gaboriau et al. [8] ultimately classified *A. polymnus* as a *Stichodactyla* specialist. However, the *A. polymnus* host classification was based on only 120 citizen science observations, and a fourth anemone species (*R. crispa*) that had been historically reported was unable to be independently verified [8]. Gaboriau et al. [8] did not define a strict set of criteria to deem a species a specialist or generalist, but one emergent result from their classification scheme was that generalists are found to have at least 3-5 hosts at the reproductive life stage, and are broadly found in association with at least 50% of all available host anemone species in their biogeographic region. Here, we revisit the underlying ecology of *A. polymnus* to test the classification of Gaboriau et al. [8] that this species is host specialist. We triple the number of host association observations by Gaboriau et al. [8], as well as integrate a comparative behavioral and stable isotope analysis of co-occurring specialist and generalist clownfishes to comprehensively characterize the ecology of *A. polymnus*. Our data demonstrate *A. polymnus* aligns with the expectations of a host generalist clownfish across multiple ecological axes. Ultimately, our findings recontextualize the results of Mercader et al. [28] on *A. polymnus* and align their findings with Gaboriau et al. [8] and Froehlich et al. [26], providing additional support for sea anemone host use being key to understanding the ecology, evolution, and phenotypic diversity of clownfishes.

## 2. Methods

### (a) <u>Amphiprion polymnus</u> host associations

Following Gaboriau et al., [8] we quantified host associations for *A. polymnus* using the citizen science platform iNaturalist. Our updated dataset incorporated n = 358 entries through 23-December-2025. We included entries with a photograph, date and GPS point, visible host anemone, and clownfish species that could be confirmed as *A. polymnus*. We discarded entries where the fish species was misidentified (n = 1) or where the host anemone was not visible (n = 50). For each entry, we identified the host species and *A. polymnus* life stage (juvenile, adult, or both present) based on their color pattern and size since clownfishes have distinct patterns with ontogenetic shifts. We removed duplicate entries of the same clownfish-anemone colony (n = 2).

### (b) Behavioral ecology of specialist and generalist

We conducted a field-based behavioral study of *A. polymnus* from 12-Jan-2025 to 24-Jan-2025 at Restorff (-5.294354, 150.104160) and Schaumann Islands (-5.293405, 150.092653), in Western New Britain, Papua New Guinea. For comparison, we also studied the behavioral ecology of one host generalist clownfish, *A. clarkii*, and one host specialist, *A. perideraion,* found on the same reef systems. Here, *A. polymnus* were found on *Stichodactyla haddoni* (n = 4), *A. perideraion* on *Radianthus magnifica* (n = 4), and *A. clarkii* on *Stichodactyla mertensii* (n = 2) and *Radianthus crispa* (n = 1). We acknowledge that our sample size is low, yet these findings replicate intraspecific behavioral and dietary patterns previously observed in Japan and Western Australia for these clownfish species [8,28].

Clownfish behaviors were filmed with GoPro12 on tripods for 2h. The first 5min were discarded to ensure divers were gone, allowing for a full 90min of video. Following Froehlich et al. [26], we measured the distance that each individual clownfish moved away from its host, a metric that captures an individual’s reliance on their host for shelter. Distance from host was measured every minute for 90 min in fish body depths, using formula (1) from Froehlich et al. [26]. We binned fish movement behavior into six distance categories based on shelter reliance: zero body depths (i.e. fish was directly in host), one body depth, 2-5 body depths, 6-10 body depths, 11-20 body depths, and 20+ body depths away. We analyzed the distance from shelter against fish species (fixed factor) and colony ID (random factor) with a categorical linear mixed model in R v4.2.2[29] with the packages ordinal[30] for analysis, rcompanion[31] for r-squared values, and tidyverse[32] for visualization.

### (c) Dietary niche of specialist and generalist

To understand the dietary niche of *A. polymnus* in comparison with co-occurring generalist (*A. clarkii*) and specialist (*A. perideraion*) clownfishes, we collected one adult from each colony filmed. Muscle tissue was sampled following euthanasia via an ice water bath. To broadly characterize multiple trophic levels, tissue samples were taken from each host anemone and reference samples from predatory fish (bonito fish and yellow spotted jack) caught by local fishermen. We pooled the anemones hosting *A. clarkii* due to low sample size for each anemone species.

On shore, all samples were dried in a dehydrator at 70 for 48h. Samples were pulverized into powder using a Bead Ruptor 12 (Omni International) if we obtained ≥10mg of sample, or by hand with metal spatulas for smaller samples. Between 0.400 to 0.600mg per sample was weighed into 3.3 x 5mm tin capsules using a microbalance. Capsules were folded into a cube, organized in 96-well plates, and sent for δ^13^C and δ^15^N analysis to the Center for Stable Isotope Biogeochemistry at University of California Berkeley. Isotopic values were provided in parts per thousand (‰) using the standard δ notation, where δX = [R_SAMPLE_/R_STANDARD_ – 1) x 1000]; with R representing high to low isotope ratios.

δ^15^N and δ^13^C values were compared across clownfish species using linear models with packages lme4[33] and LMERConvenienceFunctions[34] to generate the analysis and emmeans[35], piecewiseSEM[36] to do pairwise comparisons. Data was transformed to meet normality and homoscedasticity using Q-Q plot visualization, histograms, and residuals over fitted plots. Pairwise testing was completed using Tukey’s posthoc testing, with package emmeans. The analysis was completed within R v4.2.2[29], with additional packages: tidyverse[32], GGally[37], and reshape2[38].

### (d) Ancestral state reconstruction of host use

All underlying ecological data pointed to *A. polymnus* as a host generalist (see Results) [8]. To understand the effects of this reclassification on our understanding of the diversification of clownfishes broadly, we conducted an ancestral state reconstruction (ASR) of clownfish host-use following Gaboriau et al. [8]. We used four host categories: *Entacmaea quadricolor* specialists, *Radianthus magnifica* specialists, *Stichodactyla* specialists, and host generalists. Clownfish host categories for all species were kept consistent with Gaboriau et al. [8] except for *A. polymnus*, which we now treated as a host generalist. We used the time-calibrated species phylogenetic tree and R code from Gaboriau et al. [8] to reconstruct ancestral reproductive host association states for all clownfish species in the package corHMM [39]. We used a transition matrix with equal rates between specialists and generalists, as per Gaboriau et al. [8]. Throughout the phylogenetic tree, we generated co-occurrence between lineages with 100 stochastic maps. Additional packages were used in R v4.2.2 [29]: ape [40], mvMORPH [41], scales [42], sda [43], phytools [44], and ggtree [45].

## 3. Results and Discussion

### (a) <u>Amphiprion polymnus</u> host associations

Our new host association data for *A. polymnus* tripled the number of observations incorporated by Gaboriau et al. [8] to make their classification of *A. polymnus* as a *Stichodactyla* specialist (n = 120 vs. n = 358). Here, we recorded *A. polymnus* in association with five species of anemone hosts (versus three hosts originally [8]): *Heteractis aurora*, *Stichodactyla haddoni, Radianthus crispa, R. doreensis,* and *R. malu* (Figure 1). Our documentation of *A. polymnus* in association with *R. malu* represents a new host record for this species (Figure 1). We are also able to independently confirm that *R. crispa* served as a host for *A. polymnus* [15,21]. Raw host association frequencies (i.e. hosts to all life stages) are now 0.76 for *S. haddoni*, (Figure 1; previously 0.88 in [8]), 0.15 for *R. doreensis* (previously 0.10 in [8]), 0.04 for *R. crispa* (previously undocumented in [8]), 0.03 for *H. aurora* (previously 0.02 in [8]), and 0.02 for *R. malu* (new record). Importantly, following the reproductive adult host use paradigm established by Gaboriau et al. [8], we identify all but *R. malu* as adult habitat for *A. polymnus*. As no other reproductive host specialist clownfish species utilizes four host anemones as reproductive adult habitat [8], our updated host use data indicate *A. polymnus* is a host generalist.

**Figure 1.**
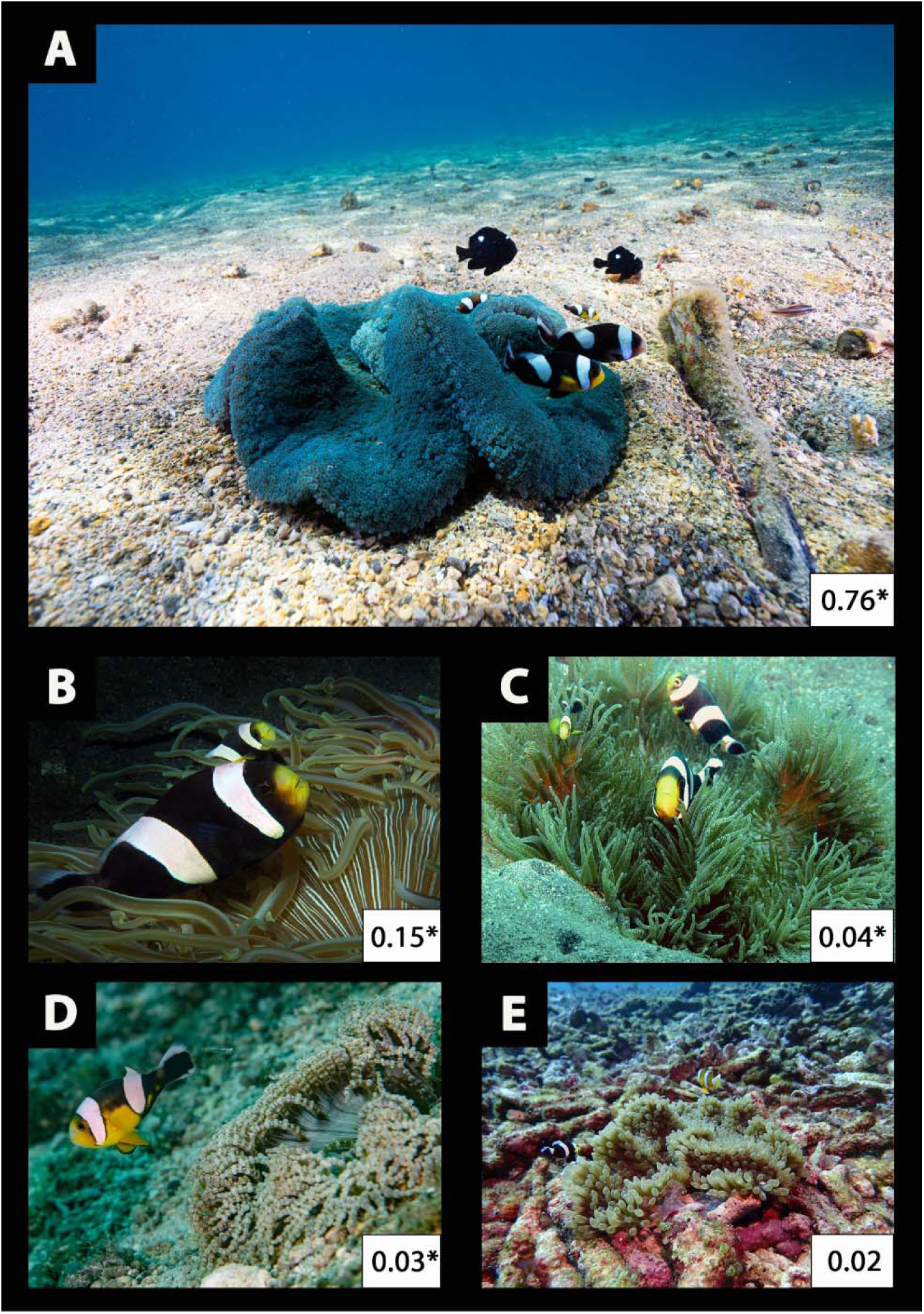
Anemone host use and association frequency of *Amphiprion polymnus*: A) *Stichodactyla haddoni,* (photo by Benjamin Titus), B) *Radianthus doreensis,* (photo by Pauline Walsh Jacobson), C) *Radianthus crispa* (photo by Bernard DuPont), D) *Heteractis aurora* (photo by Frank Krasovec), and E) *Radianthus malu* (photo by Miriam Cheng). *indicates hosts serving as reproductive adult habitat.

### (b) Behavioral ecology and dietary niche of specialist and generalist

Specialist and generalist clownfishes spent different proportions of time at different distances from their hosts (F-value = 24.74, p < 0.001, r-squared = 0.017; Figure 2). *Amphiprion polymnus* was almost never directly in their host (< 5%), occasionally near their anemone (one body depth away; ∼20%), and spent most of their time swimming and remaining far away (5+ body depths away; >75%; Figure 2A). Similarly, host generalist *A. clarkii* spent little time directly in their host (<10%), half of their time near the anemone (one body depth away; ∼55%), and the rest of the time swimming and remaining far from their host (5+ body depths away; 30%; Figure 2A). The host specialist *A. perideraion* spent most of their time in their host (∼45%) or right next to the anemone (one body depth away; ∼50%; Figure 2A).

**Figure 2.**
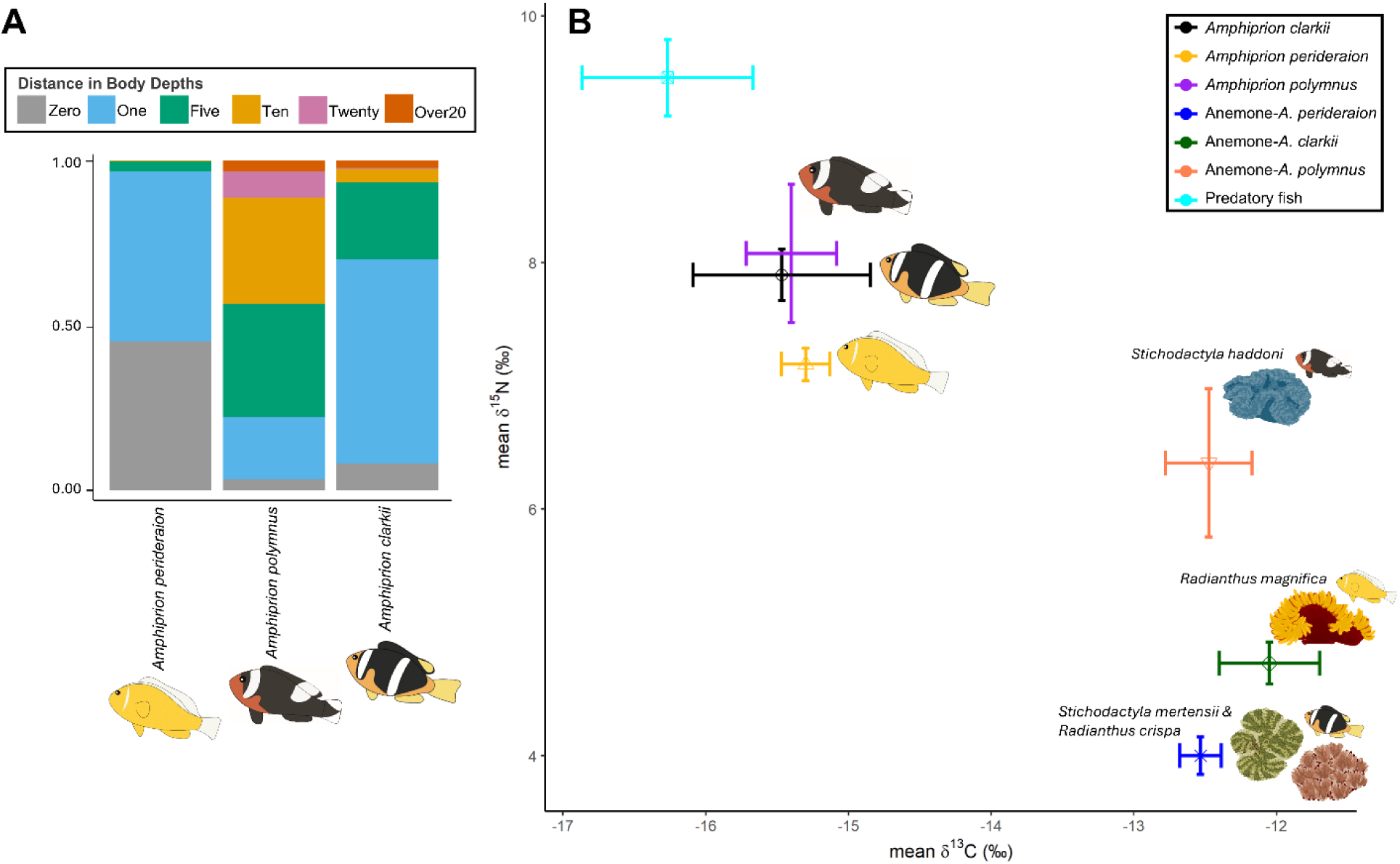
Movement behavior and dietary niche of *Amphiprion polymnus* compared to host specialist (*A. perideraion*) and host generalist (*A. clarkii*). A) Movement measured in proportion of distance that clownfish moved from their shelter, i.e. host anemone. B) Dietary niche compared to respective anemones using stable isotopes with standard error bars.

Our movement behavioral data shows that *A. polymnus* behaves like a host generalist [26], swimming and remaining far from their host. Our data represents the third independent study investigating the way *A. clarkii* and *A. perideraion* interact and rely on their host anemones for shelter, and the second study for *A. polymnus* [26,28]. These studies have been conducted across distant geographic regions and demonstrate that clownfish behaviors, and their reliance on their host, are consistent intraspecifically [26,28]. In each study, the host generalist *A. clarkii* showed little reliance on their host for shelter regardless of host identity [26,28]. Froehlich et al. [26], studied *A. clarkii* residing in *E. quadricolor* and *R. crispa* in Western Australia and added *S. mertensii* as a third host in this study. Mercader et al., [28] did not provide data on host identity but showed *A. clarkii* to have little reliance on their hosts in Japan. Similarly, *A. perideraion*, a *R. magnifica* specialist, rarely left their hosts in Western Australia [26], Japan [28], and in current study in Papua New Guinea. Specializing on *E. quadricolor*, *A. rubrocinctus* in Western Australia [26] and *A. frenatus* in Japan [28] also remained in or close to their host, as seen in *R. magnifica* and *S. mertensii* specialists, *A. ocellaris* and *A. sandaracinos* in Japan, respectively [28]. Across all of these studies a clear underlying behavioral pattern has emerged—host specialists rely heavily on their anemones and rarely travel far away; in stark contrast, host generalists consistently swim far from their hosts [26,28] (Figure 2A). Behavioral and movement data for *A. polymnus* are not significantly different than *A. clarkii* in our study in Papua New Guinea or from Mercader et al. [28] in Japan. Across both studies, *A. polymnus* behavioral ecology aligns fully with the expectations of host generalism [26].

Additional similarities between *A. polymnus* and *A. clarkii* show support for the species’ new classification as a host generalist. Mercader et al., [28] demonstrated that swimming ability, physiology, and musculature showed striking similarities between *A. clarkii* and *A. polymnus*. Our stable isotope analysis of dietary niche adds an additional ecological axis of similarity between *A. polymnus* and *A. clarkii* as they both have broader dietary niches than specialists (Figure 2B). While our mean δ^13^C values were not significantly different among our three focal clownfish species, *A. polymnus* and *A. clarkii* had significantly more δ^13^C variation than *A. perideraion* (F-value = 20.42, p < 0.001, r-squared = 0.89), but were not different from each other. *Amphiprion clarkii* and *A. polymnus* δ^15^N values were also significantly higher and more variable compared to *A. perideraion* (F-value = 16.45, p < 0.001, r-squared = 0.87), but not different from each other. The variance in dietary niche among clownfishes studied were not related to the variance in the dietary niche of their anemones (Fig 2B). These data indicate a broader diet for generalist species and narrower diet for specialists, findings consistent with data for *A. clarkii* and *A. perideraion* from Western Australia [26]. Together, the ecological and biological axes studied to date reflect a broader underlying pattern inherent to a host generalist lifestyle rather than an artifact of geography.

### (c) Ancestral state reconstruction of host use

Patterns of host use, behavior, and dietary niche all align with the conclusion that *A. polymnus* is a host generalist. Interpreted within the broader context of clownfish phenotypic diversification, our reclassification of *A. polymnus* as a host generalist, along with the results of our ancestral state reconstruction of clownfish host use (Figure 3), further strengthens the findings of Gaboriau et al. [8] that host use is driving convergent phenotypic evolution of clownfishes. Our findings reinforce those of Gaboriau et al. [8] that generalist clownfishes exhibit black base body colors, have multiple thick bars, and the common ancestor of clownfishes was likely a host generalist (Figure 3). *Amphiprion polymnus* was an outlier in their original analysis due to its black body color and thick white bars throughout ontogeny, yet was classified as a host specialist [8]. Our ancestral state reconstruction (Figure 3) now aligns patterns of generalist host use with body color for *A. polymnus* and further strengthens the conclusions [8] that reproductive host patterns are robust for broadly explaining the diversification and phenotypic evolution of clownfishes.

**Figure 3.**
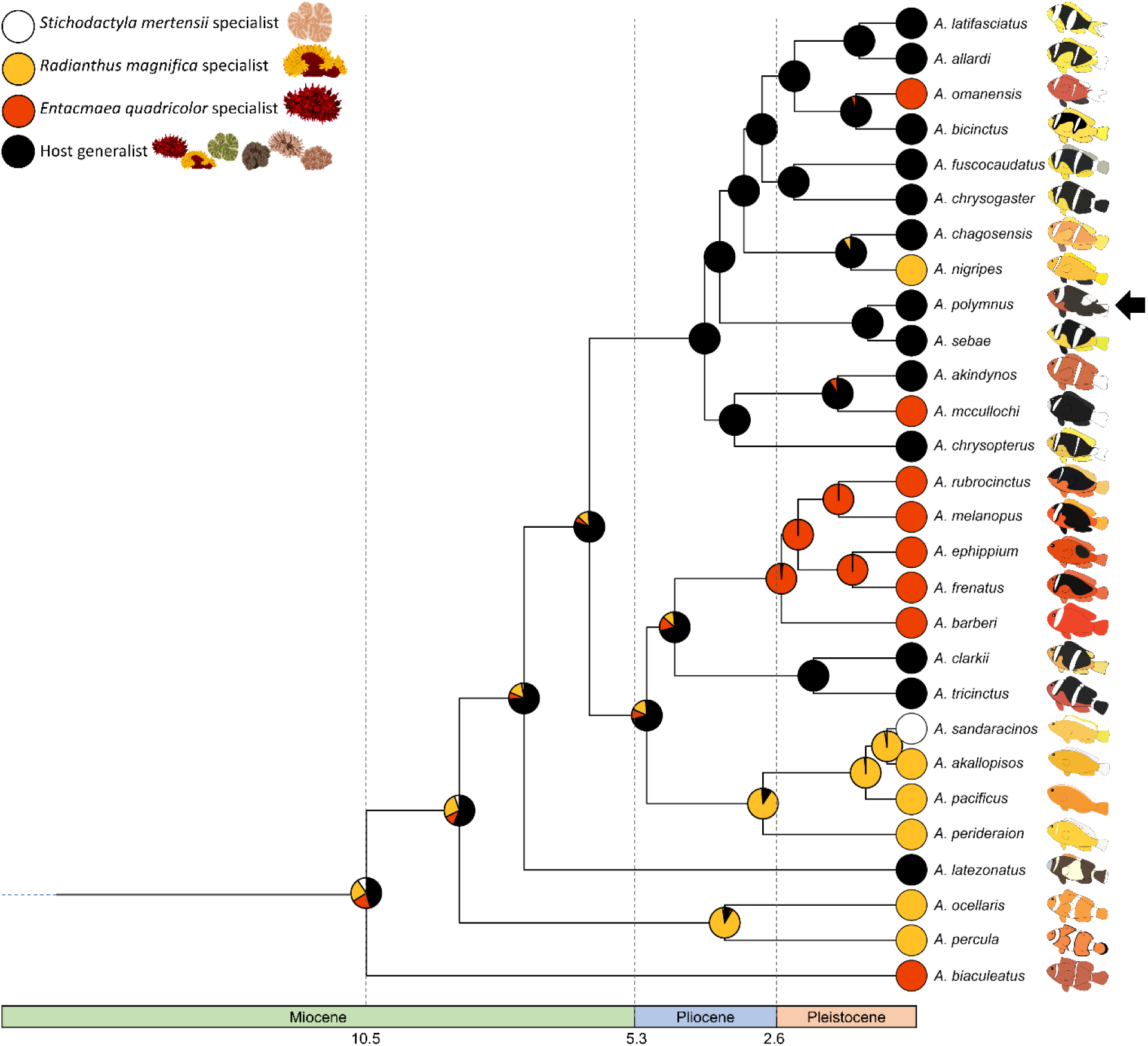
Phylogenetic tree of *Amphiprion* overlayed with updated reconstruction of ancestral host associations (internal nodes) and extant species (tip nodes). The marginal probabilities of ancestral associations for reproductive hosts are represented by pie charts at each node.

### (d) A final note on host specialism and generalism in clownfishes

In the ecological sciences, a longstanding focus has been on the many ways that species exploit available resources, develop/occupy niche space, and ultimately, co-occur [e.g. 46–51]. Species with narrow resource use, specific habitat requirements, and/or specialized functional roles are regularly defined as “specialists.” In contrast, species with broad resource use, general habitat or environmental requirements, and generalized functional roles are defined as “generalists.” This specialist-generalist paradigm has led to hundreds of theoretical and empirical studies of the processes that lead to this emergent pattern across the diverse taxa [e.g. 14,52–57]. While conceptually straightforward, nature rarely fits neatly into these discrete categories. Rather, gradients of resource use may often be encountered, which requires more nuanced organismal expertise to categorize species into specialist and generalist bins. Ultimately, the criteria used to define specialism and generalism requires careful consideration and should be regularly interrogated and refined as new data emerge.

In clownfishes, the number of host sea anemones a given fish species associates with spans a relatively continuous gradient from only a single sea anemone host to all 10 anemone hosts [8]. In the literature that precedes Gaboriau et al. [8], host specialist clownfishes have been variously defined as species that associate with one, two, or three host sea anemones [e.g. 14,17,18,58–61] yet these categorizations had relatively little ecological rationale behind these schemes other than recognizing that specialist taxa utilize a small subset of the available resources putatively available to them. Gaboriau et al. [8], in contrast, did not explicitly define criteria for specialist or generalist categories *a priori*, but rather, took a bottom-up discovery-based approach and allowed specialist and generalist categorizations to emerged from the dataset. Specifically, Gaboriau et al. [8] recognized that the frequency with which clownfishes associate with sea anemones captures the combined effects of both host preference and competition. Further, Gaboriau et al. [8] recognized that not all hosts serve as reproductive habitat for adult clownfishes and that non-reproductive hosts are developmental dead-ends rather than nursery habitats for clownfishes [62]. Taken together, natural selection will primarily act on clownfishes in their reproductive anemone hosts. This bottom-up approach led to the emergence of two categories of host specialists (e.g. red *E. quadricolor* specialists & orange *R. magnifica* specialists) and a single category of black and white hosts generalists, with one species of orange specialist *A. sandaracinos* specializing on *Stichodactyla mertensii* [8].

Following our revision of *A. polymnus* host use, we highlight the following patterns that are emerging from the revised clownfish-sea anemone interaction matrix more clearly categorize species into specialist and generalist categories (Table 1): 1) Generalists use ≥ 50% of the available sea anemone hosts found throughout their biogeographic range, whereas specialists utilize <50% of the available host sea anemones in their biogeographic range. 2) Generalists use ≥ 3 different species host sea anemones as reproductive habitat. 3) Specialists typically use only 1 host anemone as reproductive habitat, and in the occasional specialist species that uses two, the host association frequencies are heavily skewed towards just a single host (often >80% [8]).

**Table 1.**
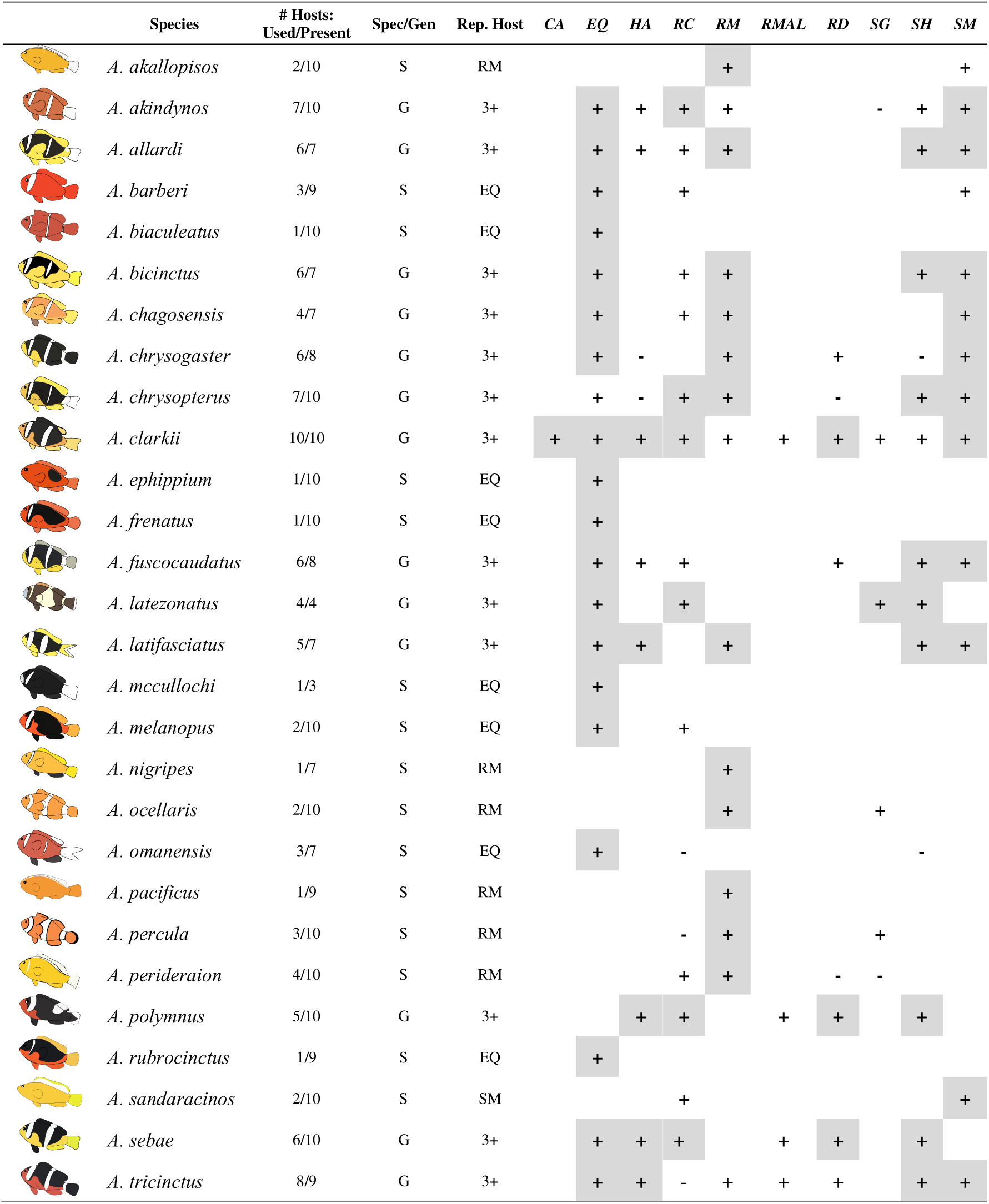
Clownfish-sea anemone association matrix. Columns from left to right include the phenotype and species name of 28 clownfishes, #Hosts: Used/Present = the number of sea anemones hosts each clownfish has been documented with compared to the number of sea anemone host species present in the fishes biogeographic range, Spec/Gen = Specialist or Generalist classification for each clownfishes, Rep. Host = the primary reproductive host for each species (note: for Generalists this is noted as 3+ sea anemone species). Sea anemone species are denoted by the following abbreviations: *CA* = *Cryptodendrum adhaesiveum, EQ* = *Entacmaea quadricolor*, *HA = Heteractis aurora, RC = Radianthus crispa, RMAG = R. magnifica, RMAL = R. malu, RD = R. doreensis, SG = Stichodactyla gigantea, SH = S. haddoni,* and *SM = S. mertensii*. A confirmed association between a clownfish species (at any life stage) and a sea anemone is denoted by (+). An association that was been previously listed by Fautin and Allen [15] but could not be independently confirmed by Gaboriau et al. [8] is listed as a (-). The primary reproductive host sea anemones for each fish are highlighted in gray. Clownfish association data were adapted from Gaboriau et al. [8] and newly collected data for *Amphiprion polymnus* from this study.

|  | Species | # Hosts:<br>Used/Present | Spec/Gen | Rep. Host | CA | EQ | HA | RC | RM | RMAL | RD | SG | SH | SM |
| --- | --- | --- | --- | --- | --- | --- | --- | --- | --- | --- | --- | --- | --- | --- |
|  | *A. akallopisos* | 2/10 | S | RM |  |  |  |  | + |  |  |  |  | + |
|  | *A. akindynos* | 7/10 | G | 3+ |  | + | + | + | + |  |  | - | + | + |
|  | *A. allardi* | 6/7 | G | 3+ |  | + | + | + | + |  |  |  | + | + |
|  | *A. barberi* | 3/9 | S | EQ |  | + |  | + |  |  |  |  |  | + |
|  | *A. biaculeatus* | 1/10 | S | EQ |  | + |  |  |  |  |  |  |  |  |
|  | *A. bicinctus* | 6/7 | G | 3+ |  | + |  | + | + |  |  |  | + | + |
|  | *A. chagosensis* | 4/7 | G | 3+ |  | + |  | + | + |  |  |  |  | + |
|  | *A. chrysogaster* | 6/8 | G | 3+ |  | + | - |  | + |  | + |  | - | + |
|  | *A. chrysopterus* | 7/10 | G | 3+ |  | + | - | + | + |  | - |  | + | + |
|  | *A. clarkii* | 10/10 | G | 3+ | + | + | + | + | + | + | + | + | + | + |
|  | *A. ephippium* | 1/10 | S | EQ |  | + |  |  |  |  |  |  |  |  |
|  | *A. frenatus* | 1/10 | S | EQ |  | + |  |  |  |  |  |  |  |  |
|  | *A. fuscocaudatus* | 6/8 | G | 3+ |  | + | + | + |  |  | + |  | + | + |
|  | *A. latezonatus* | 4/4 | G | 3+ |  | + |  | + |  |  |  | + | + |  |
|  | *A. latifasciatus* | 5/7 | G | 3+ |  | + | + |  | + |  |  |  | + | + |
|  | *A. mccullochi* | 1/3 | S | EQ |  | + |  |  |  |  |  |  |  |  |
|  | *A. melanopus* | 2/10 | S | EQ |  | + |  | + |  |  |  |  |  |  |
|  | *A. nigripes* | 1/7 | S | RM |  |  |  |  | + |  |  |  |  |  |
|  | *A. ocellaris* | 2/10 | S | RM |  |  |  |  | + |  |  | + |  |  |
|  | *A. omanensis* | 3/7 | S | EQ |  | + |  | - |  |  |  |  | - |  |
|  | *A. pacificus* | 1/9 | S | RM |  |  |  |  | + |  |  |  |  |  |
|  | *A. percula* | 3/10 | S | RM |  |  |  | - | + |  |  | + |  |  |
|  | *A. perideraion* | 4/10 | S | RM |  |  |  | + | + |  | - | - |  |  |
|  | *A. polymnus* | 5/10 | G | 3+ |  |  | + | + |  | + | + |  | + |  |
|  | *A. rubrocinctus* | 1/9 | S | EQ |  | + |  |  |  |  |  |  |  |  |
|  | *A. sandaracinos* | 2/10 | S | SM |  |  |  | + |  |  |  |  |  | + |
|  | *A. sebae* | 6/10 | G | 3+ |  | + | + | + |  | + | + |  | + |  |
|  | *A. tricinctus* | 8/9 | G | 3+ |  | + | + | - | + | + | + |  | + | + |

Finally, we want to explicitly recognize the importance of using global patterns of host associations to quantify and define specialist and generalist categories of clownfish host use. Reliance on local or regional patterns of host use may be important on those scales for understanding other processes (e.g. regional variations in color pattern of *A. clarkii*). Yet regional patterns of host use may be more reflective of the local conditions driving anemone abundance rather than global processes acting on clownfishes.

## 4. Conclusions

Our work highlights the importance of comprehensively understanding the ecology of symbioses when seeking to elucidate their role in generating biodiversity. As demonstrated with the clownfish adaptive radiation, an incomplete understanding of the underlying ecology of just one species can propel alternative hypotheses for the diversification of an entire clade. Mercader et al. [28] had concluded that host specificity was unable to explain clownfish phenotypic diversification due in part to *A. polymnus* being characterized as a host specialist albeit its movement behavior, metabolic rate, swimming performance, and musculature aligning with *A. clarkii,* a host generalist. By revisiting the ecology of *A. polymnus*, our integrative dataset demonstrates instead that the saddleback anemonefish *A. polymnus* is a host generalist across multiple ecological axes. We find direct evidence that *A. polymnus* associates with 5 anemones, is behaviorally aligned with expectations of host generalists, and also exhibits a broad and variable dietary niche like host generalists. By reinterpreting the results of Mercader at al. [28] in light of these new findings, we see striking support for characterizing *A. polymnus* as a generalist, which provides further support for reproductive host specificity driving convergent clownfish phenotypic evolution and ecology [8,26]. Integrative ecological research into additional generalist and specialist clownfish species from across *Amphiprion* will continue to be important for re-evaluating species-specific host associations for poorly known species and refining our understanding of the role of symbiotic association in driving the diversification of this iconic clade.

## Data Accessibility

The data are available at doi:10.5063/F1T72FXJ.

## Ethics Statement

The work was completed under IACUC animal ethics protocol: IACUC_22-11-6137-1.

## Author Contributions

C.F. and B.M.T.: conceptualization, funding acquisition, methodology, investigation, writing – review & editing. C.F.: project administration, data curation, formal analysis, visualization, writing – original draft.

## Competing Interests sections

There are no competing interests to report for the publication.

## Acknowledgements

We are grateful to the Restorff and Schaumann communities for granting us access to their reefs in Kimbe Bay, Papua New Guinea. We also thank Somei Jonda and the other staff members at our collaborating partner Mahonia Na Dari (https://mndpng.org/) for providing facilities, logistical support, and expert knowledge. We are grateful to captains R. Martin and B. Mautu for their incredible knowledge and for helping us navigate the reefs. We also thank all staff at Walindi Plantation Resort for their in-depth knowledge and logistical support and for providing context. We would like to thank Aurélien De Jode, Miranda Gibson, and Theresa Rueger for help in the field collection, permitting, and/or coordination.

